# Genome assembly and annotation of *Macadamia tetraphylla*

**DOI:** 10.1101/2020.03.11.987057

**Authors:** Ying-Feng Niu, Guo-Hua Li, Shu-Bang Ni, Xi-Yong He, Cheng Zheng, Zi-Yan Liu, Li-Dan Gong, Guang-Hong Kong, Jin Liu

## Abstract

Macadamia is a kind of evergreen nut trees which belong to the Proteaceae family. The two commercial macadamia species, *Macadamia integrifolia* and *M. tetraphylla*, are highly prized for their edible kernels. Catherine et al. reported *M. integrifolia* genome using NGS sequencing technology. However, the lack of a high-quality assembly for *M. tetraphylla* hinders the progress in biological research and breeding program. In this study, we report a high-quality genome sequence of *M. tetraphylla* using the Oxford Nanopore Technologies (ONT) technology. We generated an assembly of 750.54 Mb with a contig N50 length of 1.18 Mb, which is close to the size estimated by flow cytometry and k-mer analysis. Repetitive sequence represent 58.57% of the genome sequence, which is strikingly higher compared with *M. integrifolia*. A total of 31,571 protein-coding genes were annotated with an average length of 6,055 bp, of which 92.59% were functionally annotated. The genome sequence of *M. tetraphylla* will provide novel insights into the breeding of novel strains and genetic improvement of agronomic traits.

## Background & Summary

Macadamia is a kind of evergreen nut trees which belong to the Proteaceae family, *Macadamia* F. Muell. genus, grown commercially for their high-value kernels^1^. The *Macadamia* F. Muell. genus contains four different species, namely *M. integrifolia, M. tetraphylla, M. ternifolia & M. jansenii*^2^, *but only M. integrifolia, M. tetraphylla* and the hybrids of them (*Macadamia integrifolia* × *Macadamia tetraphylla*) were used to produce nuts^3-5^. Although macadamia is native to the subtropical rainforest in Queensland and New South Wales of Australia^6, 7^, but its large-scale commercial cultivation was began in Hawaii of America in 1948^8^, so it’s also known as Hawaiian Nuts.

The kernel of macadamia are rich in unsaturated fatty acids, essential amino acids, Trace elements and vitamins, the content of monounsaturated fatty acids and palmitoleic acid are also extremely rich^9-11^. Macadamia nuts can be eaten directly, also can be used as raw material for processing high-grade edible oil^12^. Because it is very nutritious, long-term consumption of macadamia helps to lower blood cholesterol, prevent arteriosclerosis, lowering the viscosity of platelets, reduce the incidence of heart disease, myocardial infarction and other cardiovascular diseases^13, 14^, therefore, the macadamia nuts were favored by consumers, known as the “Queen of nuts.”

World consumption of macadamia nuts rapid growth in recent years, according to the FAO statistics and projections, the current world macadamia total demand is more than 400,000 tons, while the supply is only about 40,000 tons, within the current and future long period of time, macadamia nuts production is still in short supply.

Macadamia commercial growing areas located mainly in tropical and subtropical areas^15^, the traditional cultivated area is the United States and Australia^16^. Because macadamia cultivation technology simple, planting high income, more cold-resistant than rubber trees, bananas and other traditional tropical crops, very suitable for some slightly cold area of tropical and subtropical countries, so the planted area growth rapidly over the past decade. However, currently the world’s macadamia orchard area in descending order are China, South Africa, Australia, Kenya, Guatemala and United States, Which China’s plantation area has reached 300,000 hectares, is the world’s largest country of Macadamia acreage.

In recent years, genome sequencing of many important tropical crops have been reported, although macadamia are diploid (2n = 28) with genome size estimates ranging from 652 Mb to 780 Mb^17-19^, but research on the genome of macadamia was very few. In 2014, 2017 and 2018, the chloroplast genome of *M. integrifolia, M. ternifolia* and *M. tetraphylla* have been sequenced by Australian and Chinese researchers^20-22^, and in 2016, the darft genome and transcriptome of *M. integrifolia* cultivar 741 have been sequenced, a total length of 518 Mb have been assembly, which spans approximately 79% of the estimated genome size^18^. But up to now, the genome sequencing of *M. tetraphylla* has not yet been reported. As an important parent species of macadamia varieties in production^23^, the genome sequencing of *M. tetraphylla* will lay a good foundation for macadamia breeding.

## Methods

### Sample collection, library construction and sequencing

A cultivated individual of *M. tetraphylla* was collected from Xishuangbanna, Yunnan Province, China. The collected plant samples were immediately frozen in liquid nitrogen and stored at -80°C before DNA isolation. High-molecular-weight genomic DNA was extracted using Qiagen plant genomic DNA extraction kit. The extracted DNA was prepared for sequencing following the manufacturer’s instructions in the genomic sequencing kit SQK-LSK108. The library was loaded on a single R9.4 flowcell and then sequenced on a GridION X5 platform. A total of 68.17 Gb data were generated with an average read length of 20.22 kb (**Table 1 and Fig. 1a**). We also generated 88.27 Gb short reads on the HiSeq2500 platform (Illumina, San Diego, CA, USA) with an insert size of 500 bp for the purpose of genome survey and assembly polishing (**Table 2**).

**Table 1.** Statistics of Oxford nanopore long-reads used for genome assembly

| Pass Reads Number | Pass Reads Bases (bp) | Mean Reads Length (bp) | Reads N50 Length (bp) | Mean Read Quality |
| --- | --- | --- | --- | --- |
| 3,382,098 | 68,173,294,275 | 20,157 | 24,256 | 7.6 |

**Table 2.**
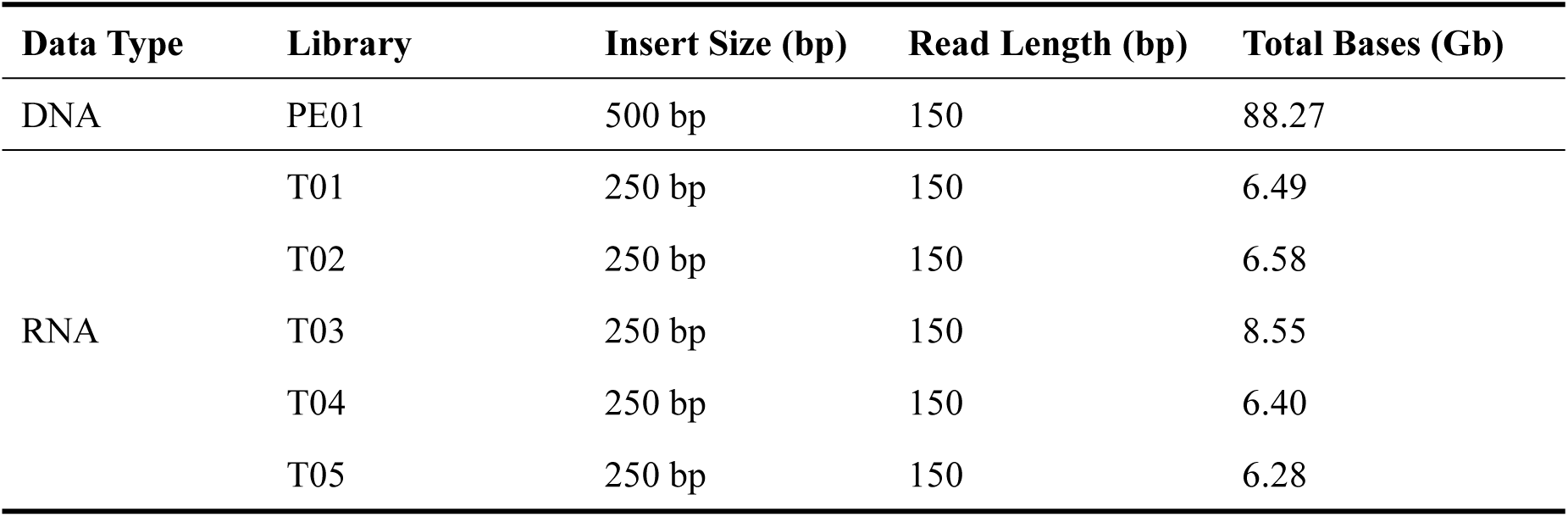
Statistics of Illumina sequencing reads used for genome and transcriptome assembly

**Figure 1.**
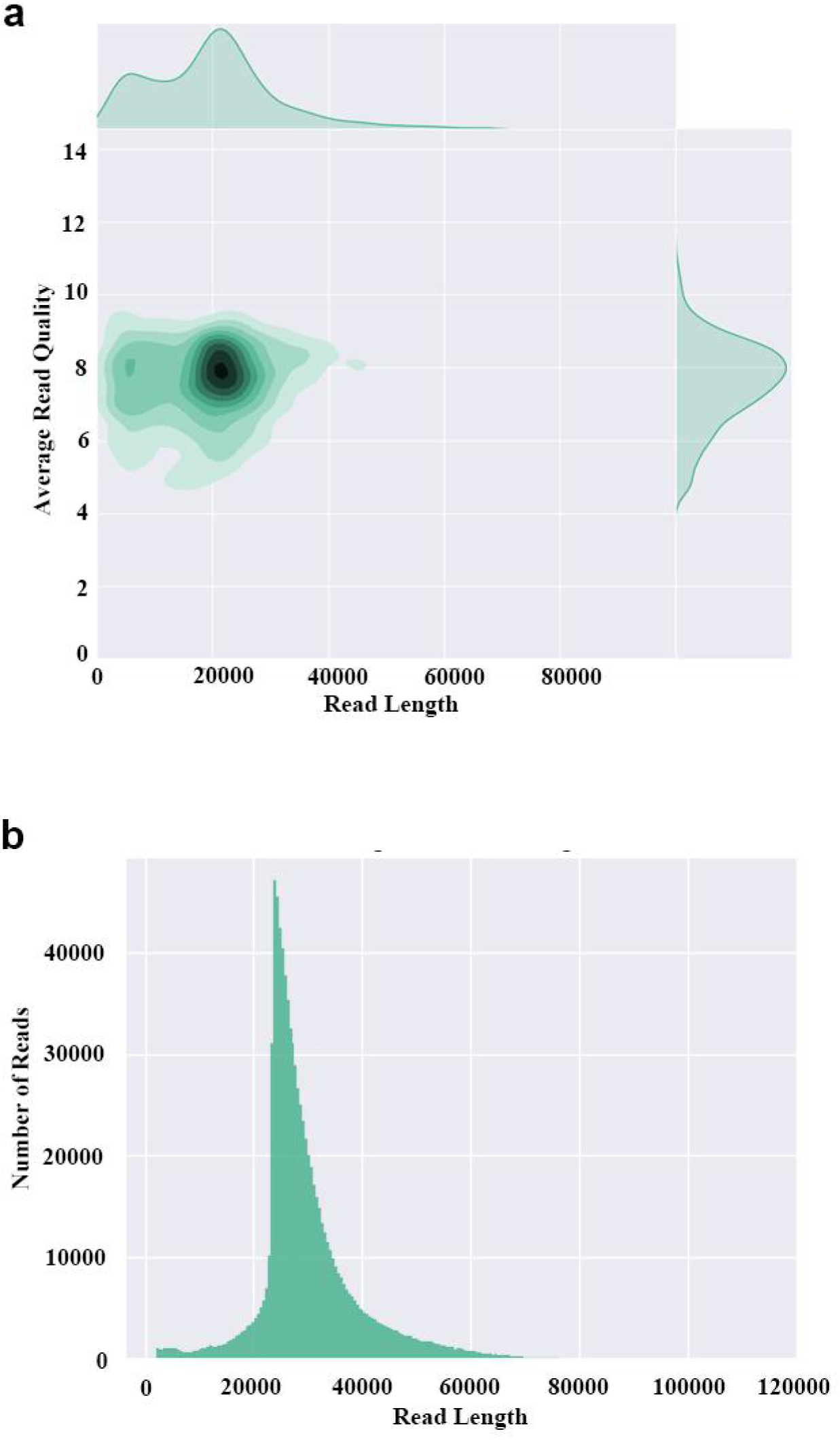
Length distribution of Oxford nanopore long reads. **(a)** Read length and read quality of the original reads; **(b)** Read length of the corrected reads.

Total RNA was isolated from five tissues (young leaves, young inflorescences, flowering inflorescence, proteoid root, and bark) using the Column Plant RNAout kit (TIANDZ, China). All cDNA libraries were prepared using an Illumina TruSeq RNA Library Prep Kit and sequenced on HiSeq2500 platform. A total of 34.3 Gb clean data were obtained after quality control (**Table 2**).

### Estimation of genome size and heterozygosity

The genome size of *M. tetraphylla* was estimated by flow cytometry following the protocol described by Dolezel^24^ and the deduced genome size was 740 Mb. We further evaluated the genome size by performing k-mer frequency analysis. In brief, Jellyfish v2.1.0^25^ was used to generate the 17-mer frequency distribution of paired end reads, the genome size was estimated according to the formula: G = K_num/peak depth (G: genome size; K_num: total number of k-mers; peak depth: depth of the major peak). The cumulative k-mer count suggested the genome size be 758 Mb (**Fig. 2**), which is similar to the result generated by flow cytometry. We estimated the heterozygosity level of the *M. tetraphylla* genome to be 1.03% using GenomeScope^26^.

**Figure 2.**
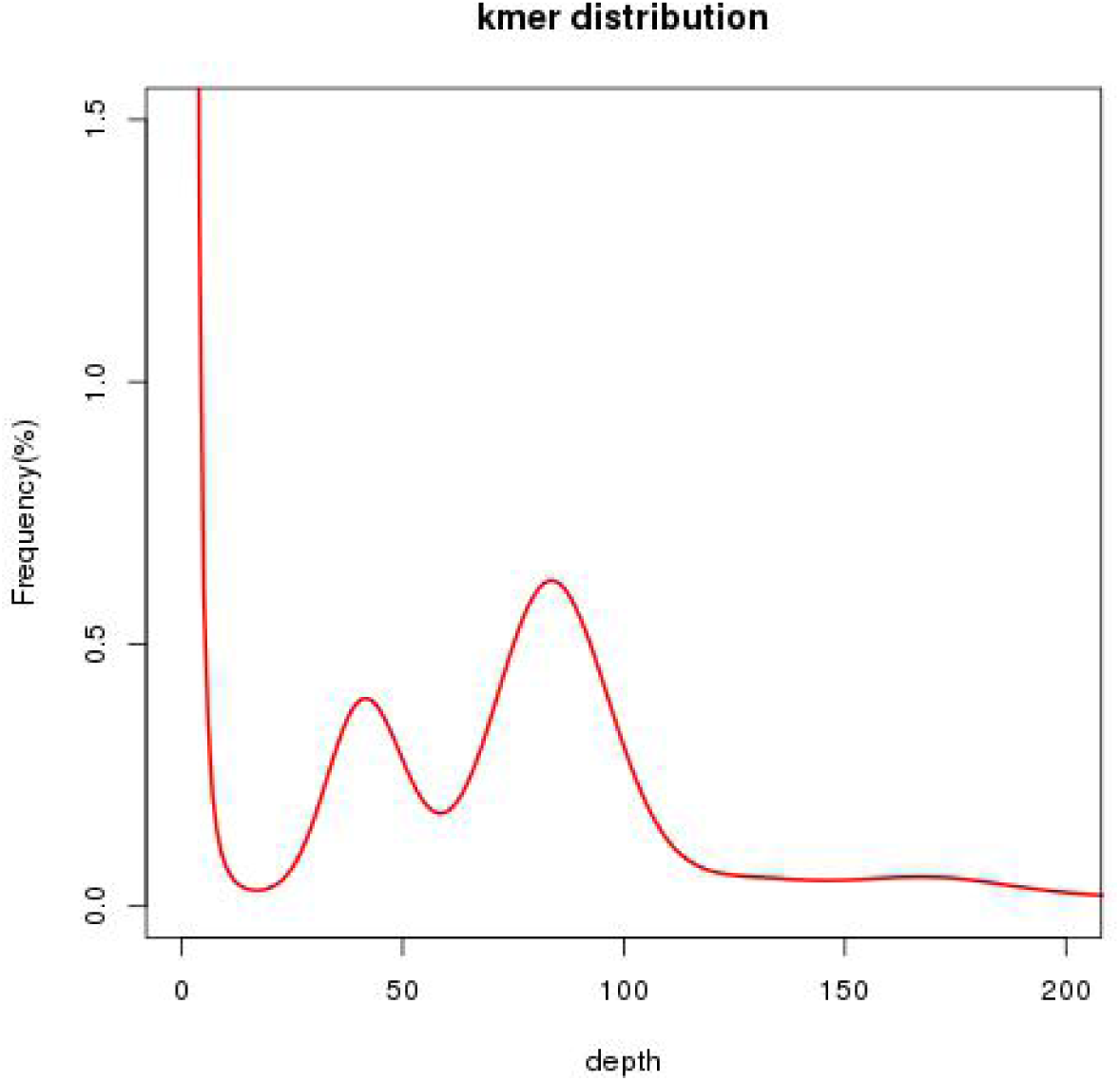
The 17-mer distribution of Illumina sequencing reads from the *M. tetraphylla* genome.

### Genome assembly

A hybrid assembly pipeline was used to alleviate the effects of the highly heterozygous genome with highly repetitive DNA sequences (**Fig. 3**). The nanopore raw reads were corrected and trimmed using Canu v1.8^27^. A total of 26.34 Gb clean data was obtained with an average length of 29.89 kb (**Fig. 1b**). The corrected reads were then fed to WTDBG v2.2^28^ for genome assembly with the following parameters: -S 2 --edge-min 2 --rescue-low-cov-edges -x ccs -g 800m. Iterative polishing was performed using Pilon v1.23^29^ to fix bases, fill gaps, and correct local misassemblies. Finally, we assembled the

**Figure 3.**
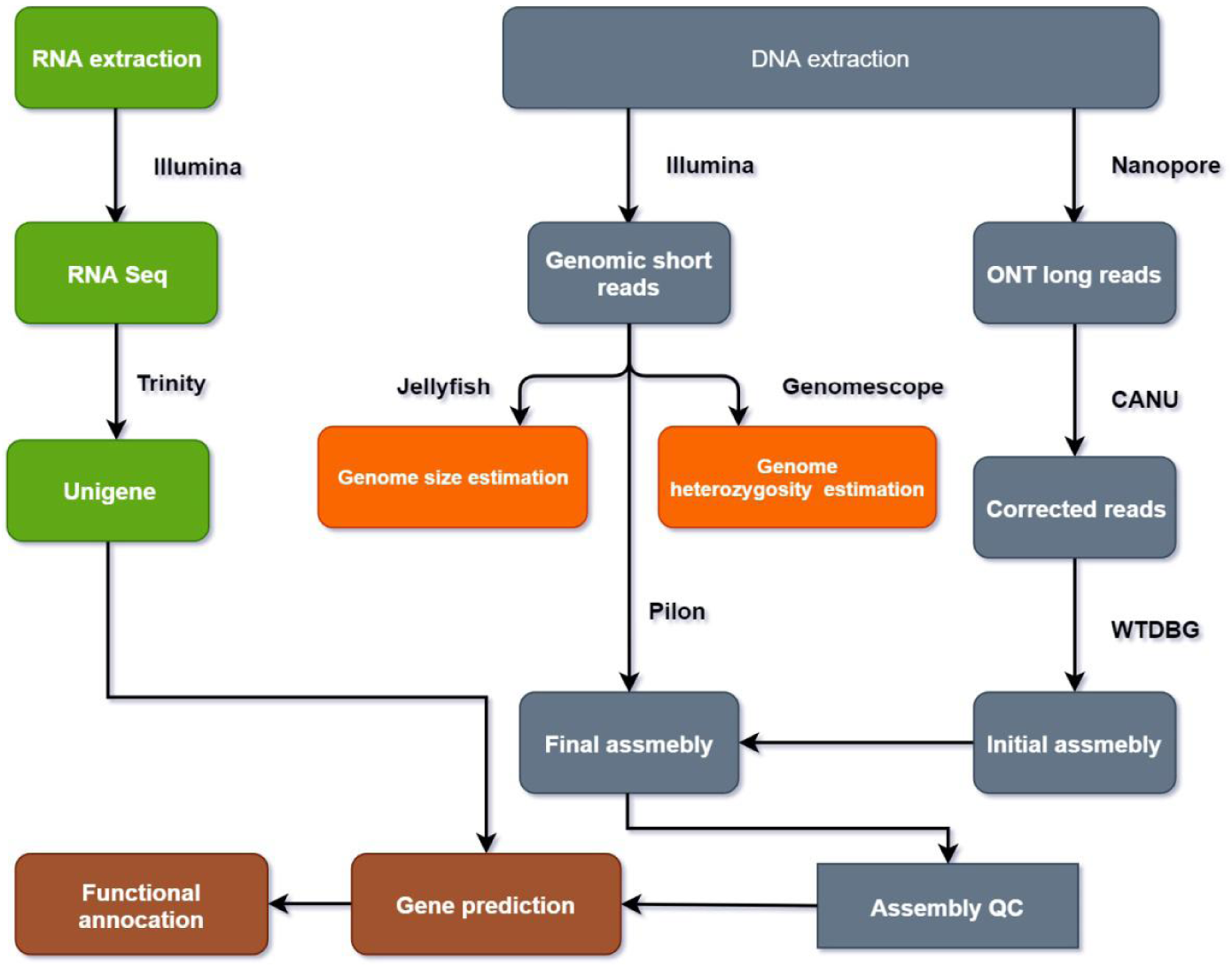
Overview of the pipeline used in this study.

*M. tetraphylla* genome into 4,335 contigs, with an N50 of 1,182,547 bp (**Table 3**). The assembly size (750 Mb) was consistent with the estimated genome size based on flow cytometry and k-mer analysis (740 Mb and 758 Mb, respectively).

**Table 3.**
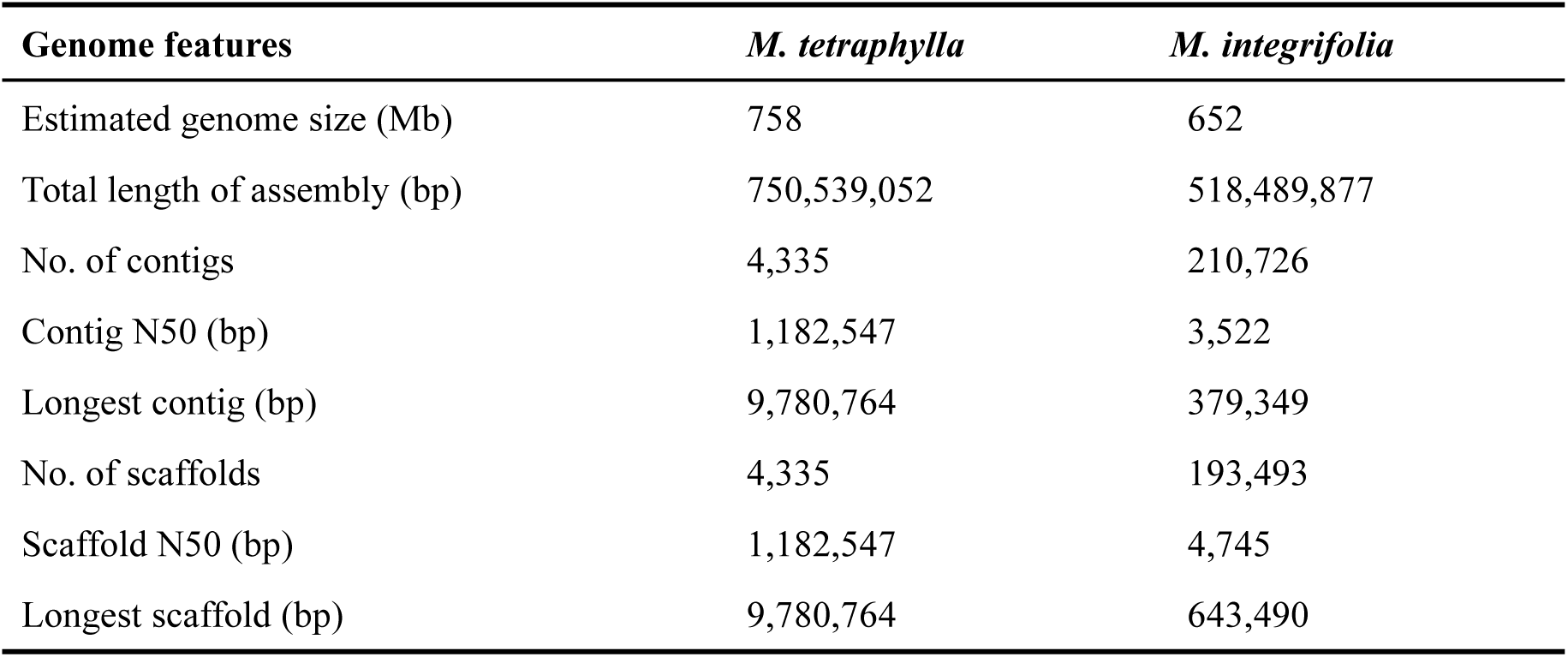
Comparison between genome assemblies for two macadamia species.

| Genome features | *M. tetraphylla* | *M. integrifolia* |
| --- | --- | --- |
| Estimated genome size (Mb) | 758 | 652 |
| Total length of assembly (bp) | 750,539,052 | 518,489,877 |
| No. of contigs | 4,335 | 210,726 |
| Contig N50 (bp) | 1,182,547 | 3,522 |
| Longest contig (bp) | 9,780,764 | 379,349 |
| No. of scaffolds | 4,335 | 193,493 |
| Scaffold N50 (bp) | 1,182,547 | 4,745 |
| Longest scaffold (bp) | 9,780,764 | 643,490 |

### Genome annotation

#### Repeat annotation

Two complementary methods were used to identity repetitive sequences in the *M. tetraphylla* genome. First, Tandem Repeats Finder v4.09^30^ was used to identity the tandem repeats. Next, a combined strategy was selected to predict transposable elements (TEs). For homology-based annotation of TEs, RepeatMasker v1.332 (http://www.repeatmasker.org) was employed to search against RepBase database (v18.07)^31^, and RepeatProteinMasker^32^ was used to search the protein database to filter TE-related proteins. Three softwares, including RepeatModeler v1.05 (http://www.repeatmasker.org/RepeatModeler.html), RepeatScout v1.05^33^, and Piler v1.06^34^, were used to construct a *de novo* library. RepeatMasker was then applied for comprehensive identification of TEs. A total of 461 Mb, which represents 61.42% of the genome (**Table 4**), were identified as repeats, much higher than that in *M. integrifolia* genome (37%). Long terminal repeat (LTR) retrotransposons represent the most predominant class of transposable elements. Result showed that *M. tetraphylla* contains 34.95% LTR retrotransposons, of which 22.00% are *Gypsy*-type elements and 5.94% are *Copia*-type elements (**Table 4**).

**Table 4.** Annotation of repeat sequences in the *M. tetraphylla* genome

| Repeat elements | Copy number | Length (bp) | Percentage (%) |
| --- | --- | --- | --- |
| Retrotransposons | 214,430 | 262,347,793 | 34.95 |
| Non-LTR retrotransposons | 72,628 | 44,629,376 | 5.95 |
| LTR retrotransposons | 141,802 | 217,718,417 | 29.01 |
| *Gypsy* | 88,052 | 165,151,480 | 22.00 |
| *Copia* | 46,649 | 44,586,669 | 5.94 |
| Other | 7,101 | 7,980,268 | 1.06 |
| DNA transposons | 11,167 | 7,521,183 | 1.00 |
| Unclassified elements | 947,252 | 191,137,916 | 25.47 |
| Total | 1,172,849 | 461,006,892 | 61.42 |

Simple sequence repeats (SSRs) were identified in the *M. tetraphylla* genome using MISA program^35^, with the following parameters: at least twelve repeats for monomer, six repeats for dimer, four repeats for trimer, three repeats for tetramer, pentamer and hexamer. A total of 510,893 SSRs were identified in *M. tetraphylla* genome (**Table 5**). Among the repeat motifs, mono-nucleotide repeats were the most predominant (**Table 5**), followed by di-, tri-, tetra-, penta- and hexa-nucleotide (**Fig. 4 and Table 5**). All these identified SSR markers may serve as potential markers of interest to *M. tetraphylla* breeding programs.

**Table 5.** Summary of simple sequence repeats (SSRs) identified in the *M. tetraphylla* genome

| Type | Unit size | Number | Length (bp) |
| --- | --- | --- | --- |
| Monomer | n >= 12 | 171,372 | 2,565,684 |
| Dimer | n >= 6 | 113,278 | 2,454,290 |
| Trimer | n >= 4 | 87,467 | 1,412,550 |
| Tetramer | n >= 3 | 81,444 | 1,079,288 |
| Pentamer | n >= 3 | 37,088 | 617,310 |
| Hexamer | n >= 3 | 20,244 | 456,330 |
| Total Number |  | 510,893 | 8,585,452 |

**Figure 4.**
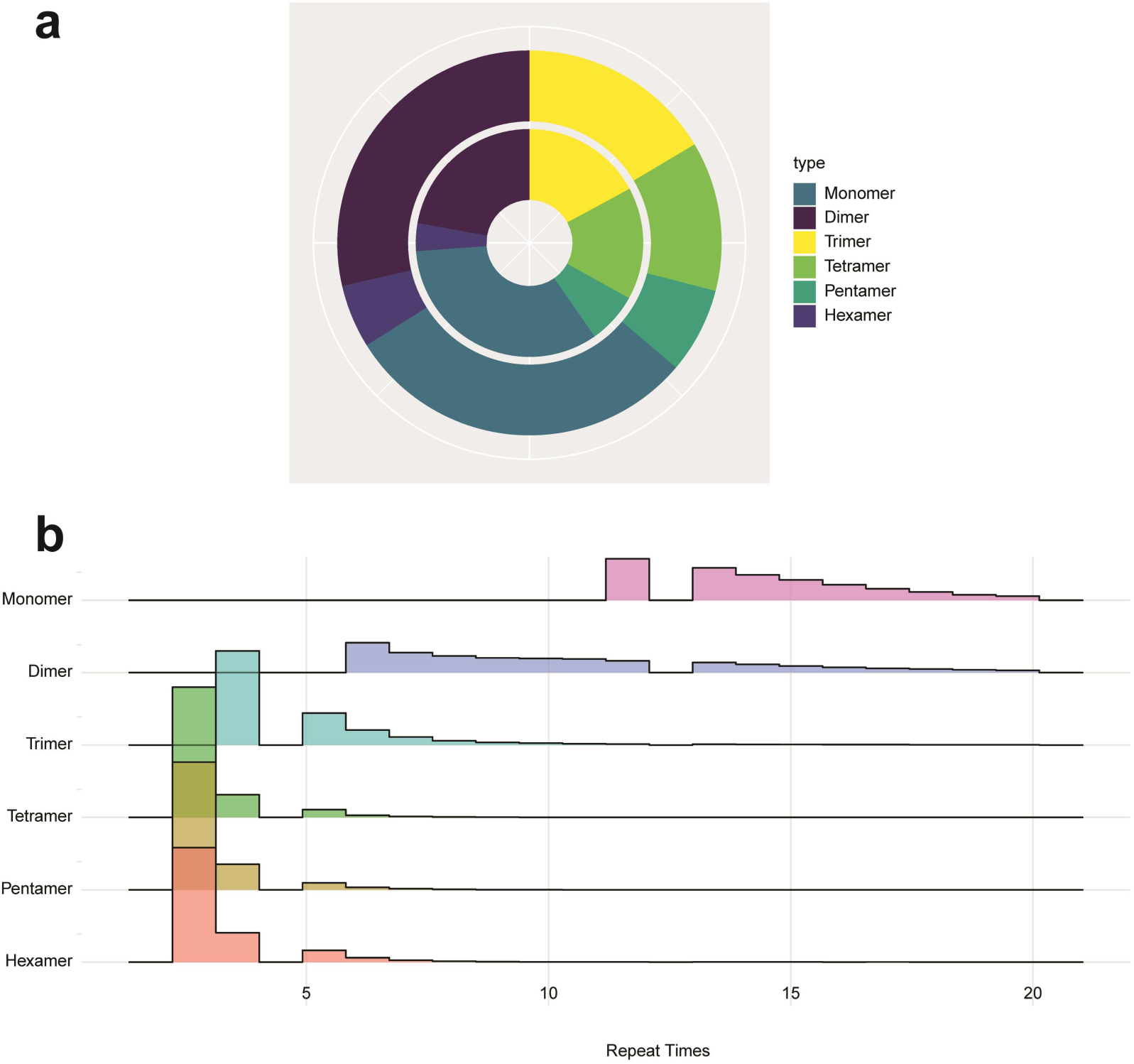
Summary of SSRs detected in *M. tetraphylla* genome. **(a**) Proportion of different types of SSRs. The outer circle represents the length proportion and the inner circle represents the number proportion; **(b)** Repeat times distribution of SSRs

#### Noncoding RNA

Five types of non-coding RNA genes, tRNA, rRNA, snRNA, snoRNA and miRNA, were identified in the *M. tetraphylla* genome. The tRNA genes were identified using tRNAscan-SE v2.0^36^. For identification of rRNA, RNAmmer v1.2^37^ was used to screen against the genome sequence. The snRNA genes were predicted using INFERNAL software (v1.1.2)^38^ with cm models from Rfam database^39^. The snoRNA genes were identified by adopting snoscan v0.9.1^40^. A total of 1,286 tRNAs, 542 rRNAs, 251 snRNAs, and 74 snoRNAs were identified in *M. tetraphylla* genome (**Table 6**).

**Table 6.** Annotation of non-coding RNA genes identified in the *M. tetraphylla* genome

| Type | Number | Average length (bp) | Total length (bp) | Percentage (%) |
| --- | --- | --- | --- | --- |
| tRNA | 1,286 | 74.34 | 95,604 | 0.0127 |
| rRNA (8s) | 210 | 112.94 | 23,718 | 0.0032 |
| rRNA (18s) | 177 | 1,886.61 | 333,931 | 0.0445 |
| rRNA (28s) | 155 | 4,333.59 | 671,707 | 0.0895 |
| snRNA | 251 | 122.01 | 30,624 | 0.0041 |
| snoRNA | 74 | 114.12 | 8,445 | 0.0011 |

#### Gene prediction and functional annotation

Gene prediction was carried out combining *de novo*, homology, and EST-based predictions. Augustus v2.7^41^ and SNAP version 2006-07-28^42^ were used to perform *de novo* prediction. The assembled transcripts generated in this study were used for iteratively self-training, and the optimized parameters were applied for further annotation by Augustus and SNAP. For homology prediction, protein sequences from *Arabidopsis thaliana*^43^, *Malus domestica*^44^, *Nelumbo nucifera*^45^, *and Rosa chinensis*^46^ were aligned to the genome using genblastA v1.0.1^47^. The hits regions were extended in both 3’ and 5’ directions and fed to GeneWise v2.2.0^48^ to obtain accurate spliced alignments. The transcripts were also mapped to the genome to generate spliced alignments using Program to Assemble Spliced Alignments (PASA) pipeline (version 2.0.2)^49^. Finally, all these predictions were consolidated into consensus gene set using EVidenceModeler (r2012-06-25)^50^. In this way, we identified 31,571 genes, with an average length of 6,055 bp and an average CDS length of 1,123 bp (**Table 7**).

**Table 7.** Prediction of protein-coding genes identified in the *M. tetraphylla* genome

| **Gene features** |  |
| --- | --- |
| Total number of protein-coding genes | 31,571 |
| Gene length in genome (%) | 25.47 |
| Average gene length (bp) | 6,055 |
| Average exon length (bp) | 222 |
| Average CDS length (bp) | 1,123 |
| Average intron length (bp) | 1,213 |
| Average exons per gene | 5.1 |
| Gene density (gene per Mb) | 42.04 |

Functional assignment was carried out by performing blastp searches (with e value 1e-5) against SwissProt database^51^. The KAAS server^52^ was used to map the predicted genes onto KEGG metabolic pathways. InterProScan v5.10–50.0^53^ determined the motifs and functional domain. The GO term and pfam domain were directly obtained from the InterProScan results. A total of 29,233 genes (92.59%) were functionally annotated by these methods (**Fig. 5 and Table 8**).

**Table 8.** Functional annotation of the protein-coding genes

| **Database** | **Number** | **Percentage (%)** |
| --- | --- | --- |
| **Swiss-Prot** | 22,869 | 72.44 |
| **KEGG** | 8,303 | 26.30 |
| **InterPro** | 29,052 | 92.02 |
| **GO** | 17,864 | 56.58 |
| **Pfam** | 21,925 | 69.45 |
| **Total** | 29,233 | 92.59 |

**Figure 5.**
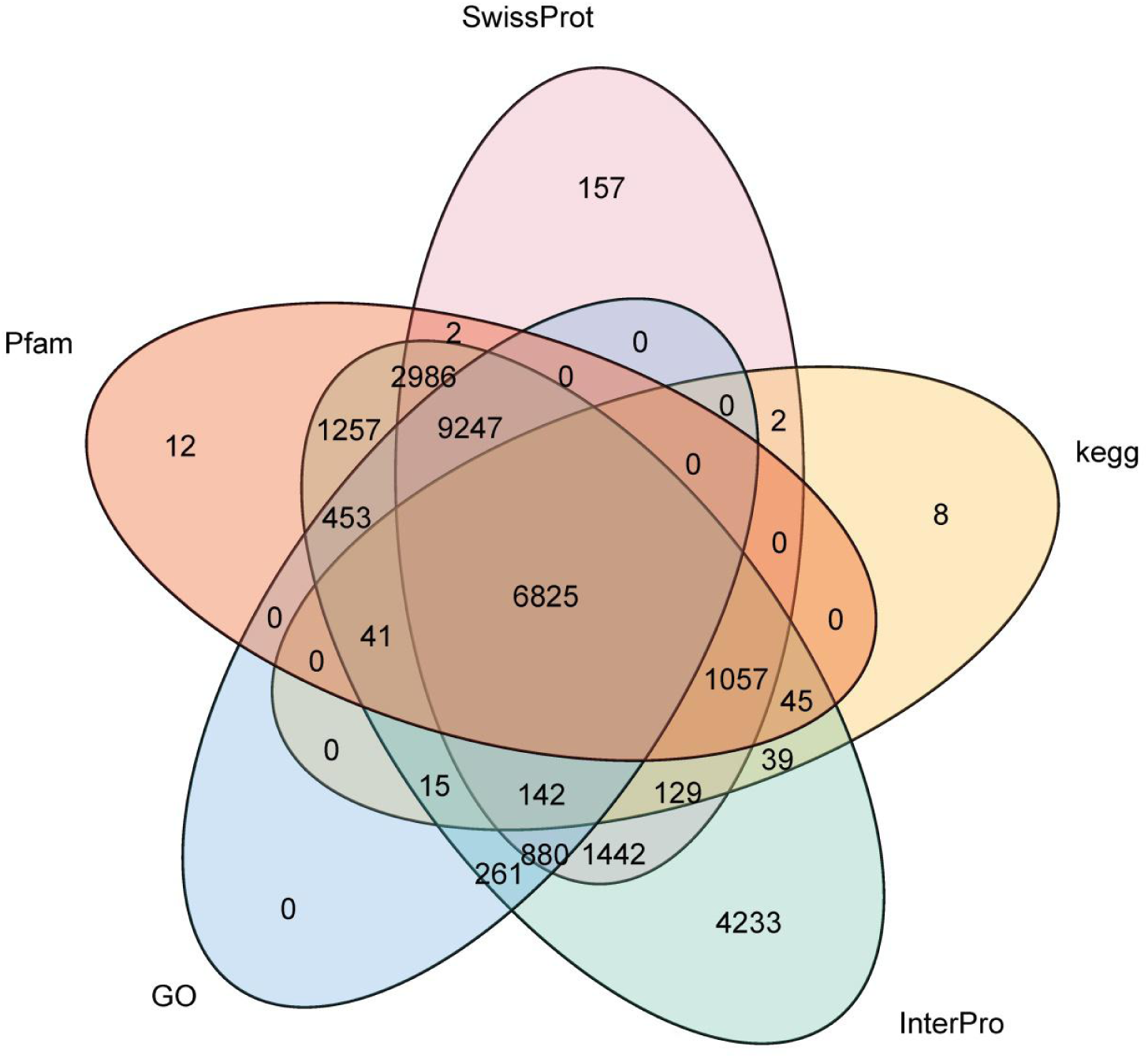
Venn diagram of functional annotation.

#### Data Records

All sequencing Illumina genomic reads and Oxford nanopore long reads have been deposited in the NCBI Sequence Read Archive (SRA) under the accession number SAMN13219572. RNA-seq reads of five tissues were also available at NCBI SRA database with the accession number SAMN13241873, SAMN13241875, SAMN13241876, SAMN13241877 and SAMN13241873.

### Technical Validation

#### Quality filtering of Illumina short reads

The Illumina raw reads were trimmed using Trimmomatic v0.32^54^ for quality. The adaptors were removed, low-quality and N base reads were trimmed. The reads were cut when the average quality per base drops below 15 with a 4-base wide sliding window.

#### Evaluation of genome assembly

To assess the completeness and accuracy of the genome assembly, the Illumina sequencing reads were mapped to the genome using bowtie2 v2.2.6^55^. Result showed that 94.25% of the short reads could be mapped to the genome, with an 87.84% properly paired mapping rate (**Table 9**). Additionally, the assembly was evaluated by BUSCO (Benchmarking Universal SingleCopy Orthologs)^56^. About 89.72% (1,292 out of the 1,440) conserved genes in embryophyta lineage were present in the assembly (**Table 9**). To further evaluate the genome assembly, the RNA reads were mapped to the genome using HISAT2^57^ with a 92.00% success rate (**Table 9**). We have also calculated the GC content with a 2kb non-overlapping sliding window and there is no obvious GC bias in the genome assembly (**Fig. 6**). All these results suggested a high quality of *M. tetraphylla* genome.

**Table 9.**
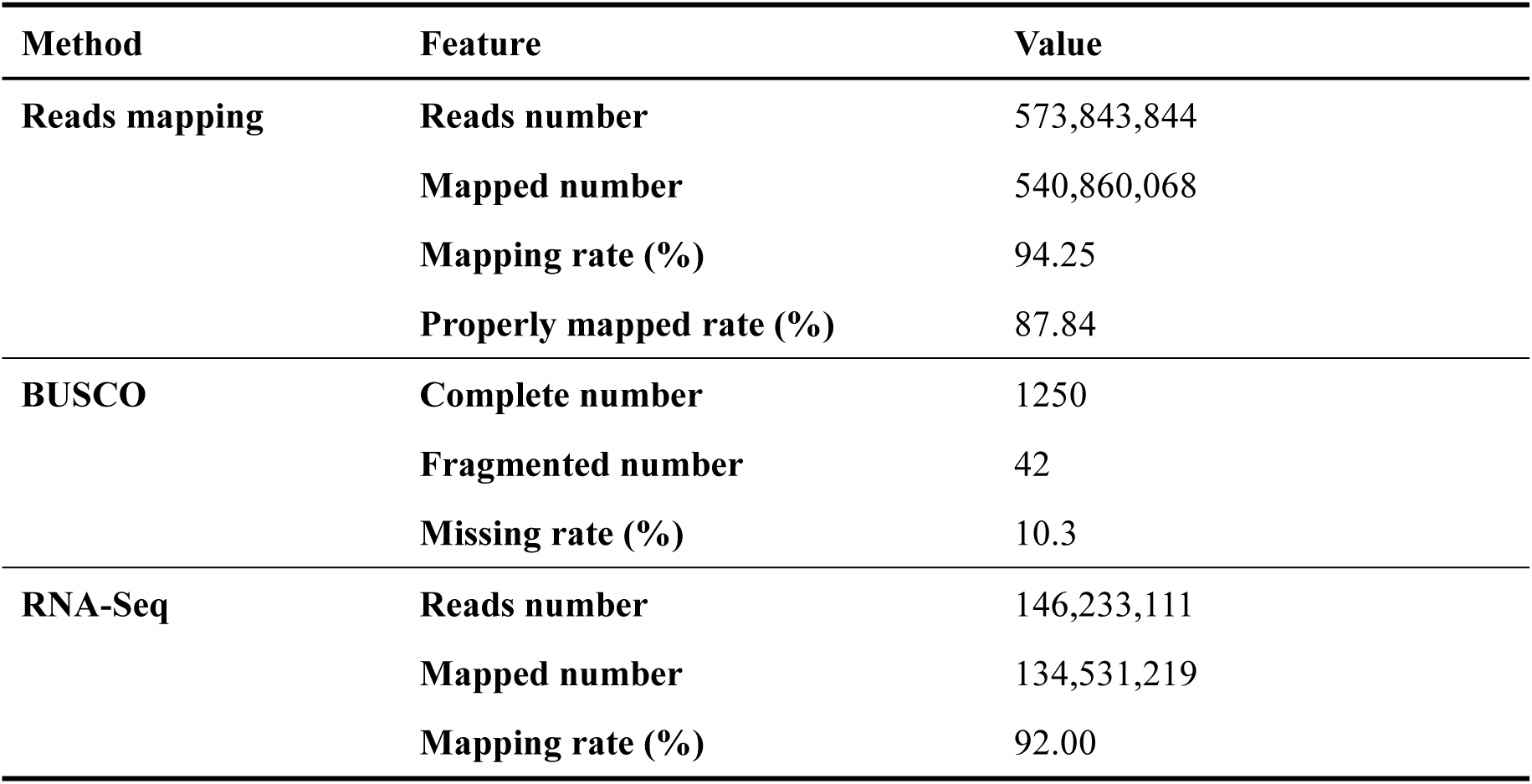
Evaluation of genome assembly using different methods

| Method | Feature | Value |
| --- | --- | --- |
| **Reads mapping** | **Reads number** | 573,843,844 |
|  | **Mapped number** | 540,860,068 |
|  | **Mapping rate (%)** | 94.25 |
|  | **Properly mapped rate (%)** | 87.84 |
| **BUSCO** | **Complete number** | 1250 |
|  | **Fragmented number** | 42 |
|  | **Missing rate (%)** | 10.3 |
| **RNA-Seq** | **Reads number** | 146,233,111 |
|  | **Mapped number** | 134,531,219 |
|  | **Mapping rate (%)** | 92.00 |

**Figure 6.**
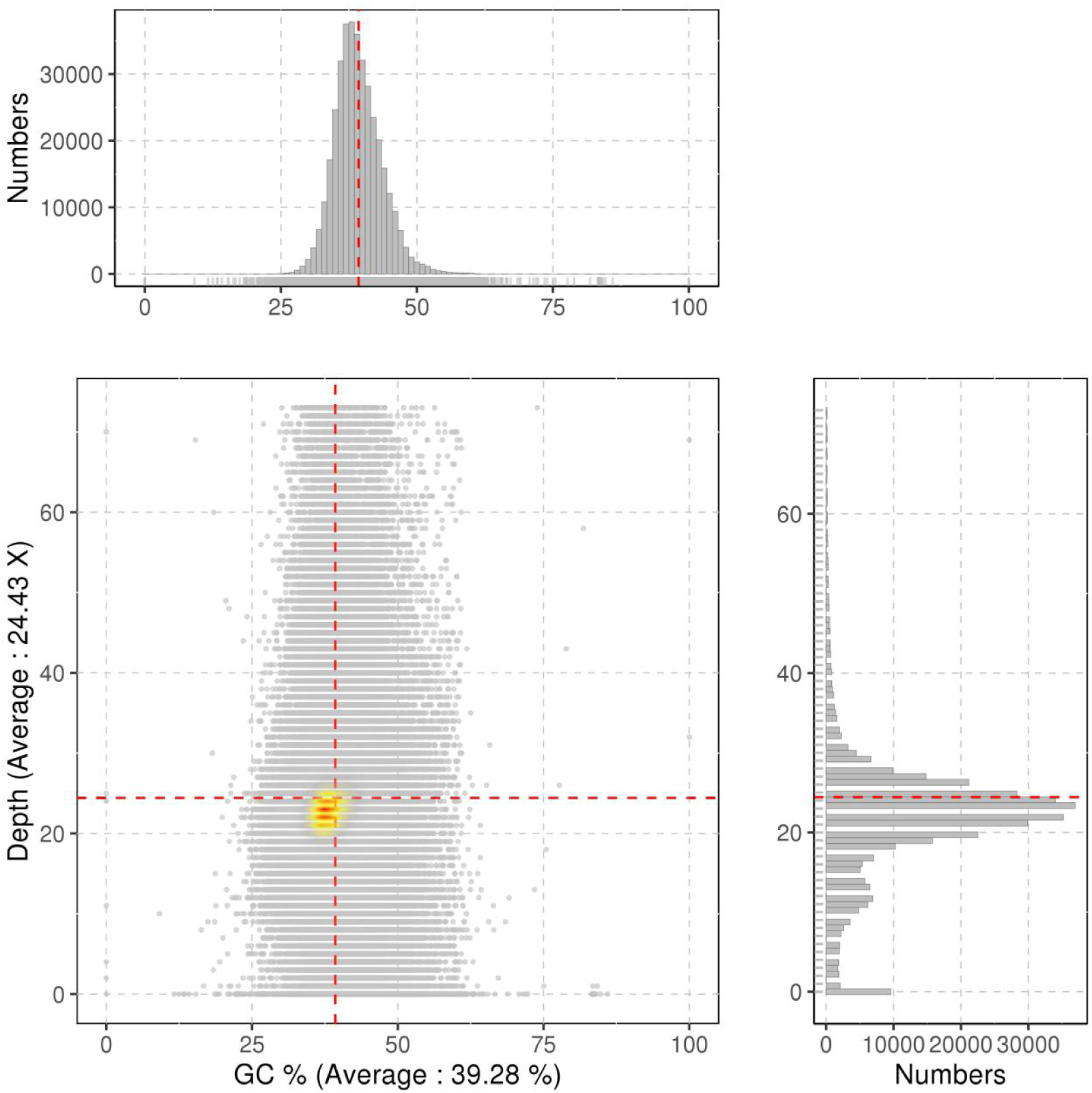
The GC content distribution of the *M. tetraphylla* genomes.

### Code Availability

The execution of this work involved many software tools, whose versions, settings and parameters are described below. (1) **Jellyfish:** version 2.1.0, -m = 17; (2) **GenomeScope:** -k = 17; (3) **Canu:** version 1.8, correctedErrorRate=0.050; (4) **WTDBG:** version 2.2, -S 2 --edge-min 2 --rescue-low-cov-edges -x ccs -g 800m; (5) **Pilon:** version 1.23, default parameters; (6) **Tandem Repeats Finder:** version 4.09, default parameters; (7) **RepeatMasker:** version 1.332, e ncbi -pa 20; (8) **RepeatProteinMasker:** default parameters; (9) **RepeatModeler:** version 1.05, -engine ncbi -pa 20; (10) **RepeatScout:** version 1.05, default parameters; (11) **Piler:** version 1.06, default parameters; (12) **MISA:** 1-12 2-6 3-4 4-3 5-3 6-3; (13) **tRNAscan-SE:** version 2.0, default parameters; (14) **RNAmmer:** version 1.2, -S euk -m lsu,ssu,tsu -f rRNA.fasta -gff rRNA.gff2 -h rRNA.hmmreport; (15) **INFERNAL:** version 1.1.2, default parameters; (16) **snoscan:** version 0.9.1, default parameters; (17) **Augustus:** version 2.7, --gff3=on; (18) **SNAP:** version 2006-07-28, default parameters; (19) **genblastA:** version 1.0.1, -e 1e-5 -c 0.5 -d 60000; (20) **GeneWise:** version 2.2.0, default parameters; (21) **PASA:** version 2.0.2, default parameters; (22) **EVidenceModeler:** version r2012-06-25, default parameters; (23) **blastp:** version 2.2.26, default parameters; (24) **InterProScan:** version v5.10–50.0, -goterms; (25) **Trimmomatic:** version 0.32, LEADING:3 TRAILING:3 SLIDINGWINDOW:4:15 MINLEN:36; (26) **bowtie2:** version 2.2.6, default parameters; (27) **Trinity:** version 2.8.4, --seqType fq --max_memory 600G --no_normalize_reads --full_cleanup --min_contig_length 250; (28) **HISAT2:** default parameters.

## Acknowledgements

This work was supported by the National Natural Science Foundation of China (No. 31760215), the Technology Innovation Talents Project of Yunnan Province (2018HB086), and Sci-Tech Innovation System Construction for Tropical Crops Grant of Yunnan Province (RF2019/RF2020).

## Author Contributions

Y.-F.N. and J.L. conceived the study and managed the project. G.-H.L., S.-B.N., and X.-Y.H. designed the scientific objectives. C.Z., Z.-Y.L., L.-D.G., and G.-H. K.collected the samples and extracted the genomic DNA. J.L. estimated the genome size, assembled the genome and carried out the gene annotation. Y.-F.N. wrote the manuscript, and all authors contributed to writing and editing the final manuscript.

## Additional Information

